# Aligned representation of visual and tactile motion directions in hMT+/V5 and fronto-parietal regions

**DOI:** 10.1101/2025.03.24.645080

**Authors:** Iqra Shahzad, Ceren Battal, Filippo Cerpelloni, Alice Van Audenhaege, André Mouraux, Olivier Collignon

**Author notes:** Correspondence should be addressed to: Iqra Shahzad and Olivier Collignon.

## Abstract

Moving events on the skin can be perceived through vision and touch. How does the brain create a unified multisensory representation of motion directions initially acquired in different coordinate systems? We show that the middle occipito-temporal region (hMT+/V5), along with a fronto-parietal network, encodes visual and tactile directions using a common external frame of reference independent of body posture. We characterized brain activity using fMRI in participants exposed to directional visual and tactile motion stimuli across different hand postures. We demonstrate that individually and functionally defined hMT+/V5 shows univariate preference for both visual and tactile motion and encodes motion directions across hand postures. Unlike somatosensory regions, information about tactile directions was enhanced in right hMT+/V5 when mapped using an external as compared to a somatotopic frame of reference. Crossmodal decoding showed that tactile directions defined in an external frame of reference, but not a somatotopic one, align with the representation of visual directions in the right hMT+/V5 (both in MT and MST). A whole brain searchlight group analysis confirmed these individually defined regions-of-interest results and extended the presence of an aligned visual-tactile code for directional motion in external space to the parietal and frontal cortex. Our findings reveal a brain network involving hMT+/V5 that encodes motion directions in vision and touch using a common external frame of reference.

## Introduction

Imagine a mosquito landing on your arm—you instinctively move to swat it. Behind this seemingly simple action lies a remarkable computational challenge for the brain: it must integrate directional motion information from multiple sensory modalities, such as vision and touch. These computations are especially complex because the sensory signals are initially encoded in distinct reference frames—retinotopic for vision and somatotopic for touch - while the sensory organs themselves (eyes and limbs) are constantly moving through space (Heed et al., 2015). To overcome this challenge, the brain must engage in a reformatting process - either remapping signals from one modality into the other’s reference frame or transforming both into a shared, external frame that transcends their original topographic organization (Avillac et al., 2004, 2005). Behavioral evidence from bidirectional crossmodal motion aftereffects - where motion adaptation transfers between vision and touch—suggests that motion processing indeed relies on shared representations across modalities (Konkle et al., 2009). Yet a crucial question remains: where in the brain are these aligned motion representations instantiated?

In primates the alignment of multisensory spatial information has traditionally been attributed to multisensory convergence zones such as the parietal and frontal cortices (Meyer & Damasio, 2009) (Bremmer et al., 2001; Rizzolatti et al., 1981) (Avillac et al., 2004; Duhamel et al., 1997, 1998). Within the parietal cortex, neurons in the Ventral Intraparietal (VIP) area exhibit preference for both visual (Colby et al., 1993) and tactile motion signals (Duhamel et al., 1998; Summers et al., 2009), with partially aligned receptive fields (RFs) across modalities (Avillac et al., 2005). Similarly, premotor areas contribute to transforming tactile inputs from a somatotopic to an external spatial representation (Crollen, Lazzouni, et al., 2017; Lloyd et al., 2003; Matelli & Luppino, 2001). Beyond these fronto-parietal regions, the human middle temporal complex (hMT+/V5) - long established as a critical node for visual motion processing (Albright, 1984; Beckers & Zeki, 1995; Zeki et al., 1991; Zihl et al., 1983), also shows selective response to moving sounds (Battal et al., 2022; Dormal et al., 2016; Poirier et al., 2005; Rezk et al., 2020) and to tactile motion ((Blake et al., 2004; Hagen et al., 2002; Ricciardi et al., 2007); but see (Jiang et al., 2015)). Moreover, hMT+/V5 encodes visual, auditory and tactile motion directions (Rezk et al., 2020)(van Kemenade et al., 2014) and is causally involved in tactile motion direction discrimination (Amemiya et al., 2017; Ricciardi et al., 2011). However, whether hMT+/V5 transforms incoming moving signals across the senses into a unified frame of reference independent of body posture remains unknown. Previous studies have shown that hMT+/V5 responses to visual motion are modulated by gaze direction, suggesting a non-retinotopic (“spatiotopic”) coding format (Burr & Morrone, 2011; Crespi et al., 2011; d’Avossa et al., 2007; Melcher & Morrone, 2003) - a mechanism that could also support multisensory integration of motion signals.

In this study, we investigated whether hMT+/V5 along with fronto-parietal regions encode visual and tactile motion directions using a shared neural code that is independent of body posture. In a first functional Magnetic Resonance Imaging (fMRI) experiment, we used independent visual and tactile motion localizers to individually and functionally define motion selective regions across both modalities. In a second fMRI experiment, we presented directional motion in both vision and touch while manipulating hand posture to systematically alter the perceived axis of tactile motion directions. Using multivariate pattern analyses (MVPA) within individually defined functional regions of interest (fROIs), complemented by whole-brain searchlight analyses, we reveal an occipito-parieto-frontal network, including hMT+/V5, that represents visual and tactile motion using an aligned format independent of body posture.

## Methods

### Participants

Twenty healthy participants (11 females; mean age ± SD = 25.15 ± 3.51 years; age range = 19–33 years) were recruited for this study. All participants were right-handed, as determined using the Edinburgh Handedness Inventory (Oldfield, 1971) and had normal or corrected-to-normal vision. Two participants volunteered exclusively for the directional motion decoding experiments but did not participate in the localizer experiments. Consequently, the localizer experiments were conducted with only eighteen participants. All participants were naïve to the purpose of the study and reported no history of neurological or psychiatric disorders. The study protocol was approved by the Research Ethics Committee of Cliniques Universitaires Saint-Luc and Université Catholique de Louvain (UCLouvain). Written informed consent was obtained from all participants before the experiments, and they received monetary compensation for their participation. All the experiments were conducted over a minimum of two sessions, each lasting approximately 1 hour and 45 minutes.

### Experimental Setup

The experiments were conducted using a 3-Tesla GE MR scanner equipped with a 48-channel head coil (Signa™ Premier, General Electric Company, USA; serial number: 000000210036MR03; software version: 28-LX-MR, release: RX28.0_R04_2020.b). Participants lay comfortably on the scanner bed, viewing a screen positioned behind the scanner. Throughout the experiment, they held a response button in their left hand. For the tactile motion localizer, participants kept their right hand open (palm up) besides their body to facilitate tactile stimulation. In the directional motion decoding experiments, participants were instructed to adopt one of the two hand positions depending on the experimental condition. In the hand-up (HU) position, their forearm was positioned perpendicular to the body plane with the palm facing outward, while in the hand-down (HD) position, the forearm was kept parallel to the body plane, with the palm also facing outward (Figure 1A).

**Figure 1.**
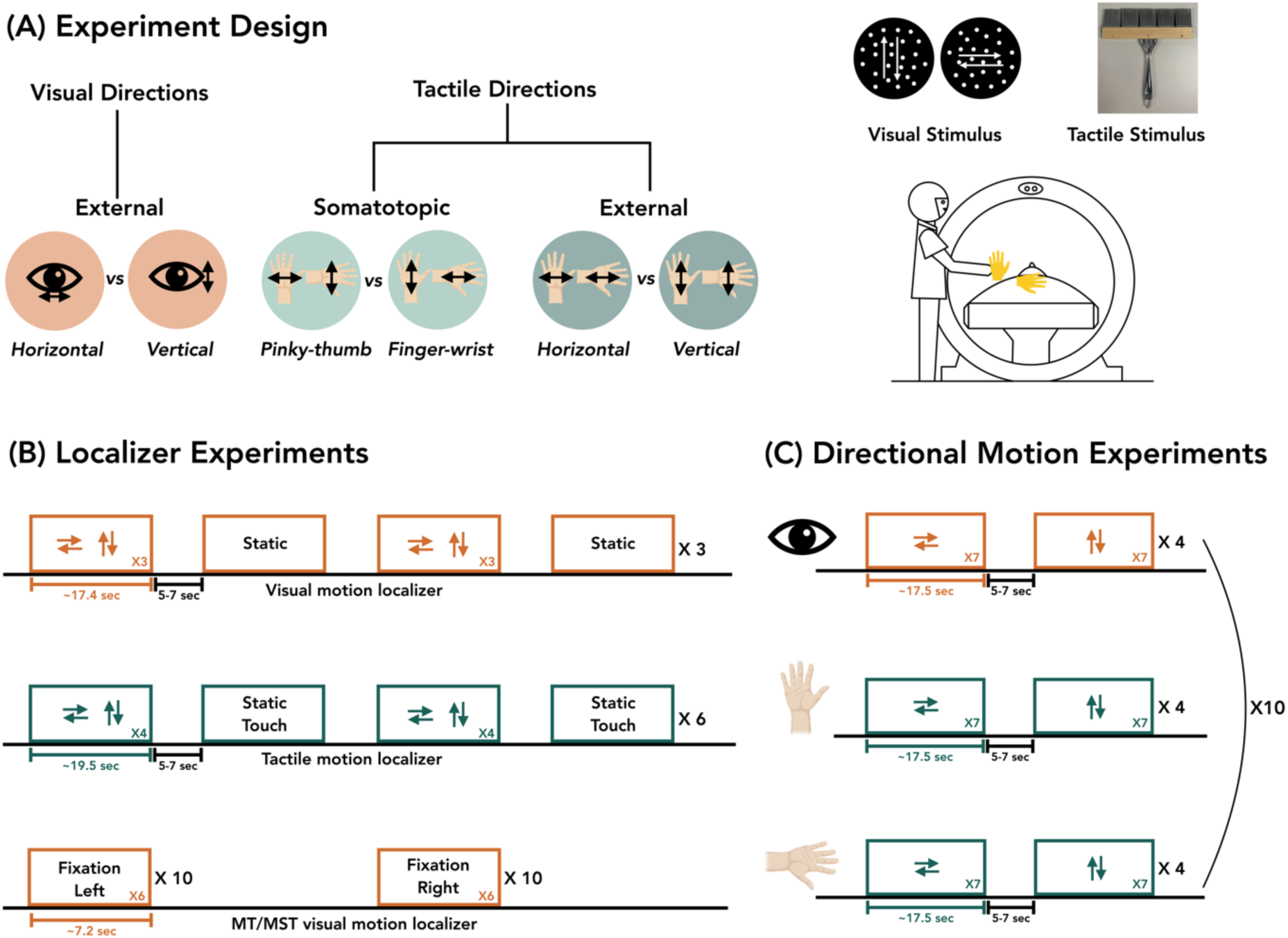
**Experiments.** (A) Experiment design. 2 tactile frames of reference X 2 tactile motion directions X 2 visual motion directions. The tactile frames of reference were manipulated using two hand positions - Hand Up and Hand Down. Within each hand position, the tactile stimulation is either along the axis of pinky-thumb or finger-wrist. Thus, with the manipulation of hand postures, the tactile motion directions were either defined somatotopically (pinky-thumb and finger-wrist) or in the external space (horizontal and vertical). The two visual axis-of-motion directions defined in the external space were horizontal and vertical. (B) Localizer Experiments. Experiments were performed to functionally and individually define the Regions-of-Interest (ROIs). The visual and tactile motion localizers consisting of alternating motion and static stimuli were used to functionally identify visual and tactile motion selective regions respectively. MT/MST visual motion localizer consisting of visual radial motion (approaching and receding) in alternating left and right hemifield was performed to identify MST and MT areas in each individual. (C) Directional Motion Decoding Experiments. 30 runs were performed with each participant, where each run consisted of only directional motion stimulation. 10 runs corresponded to a single sensory modality and one hand position. One beta map corresponding to each motion direction was obtained from 1 run. Thus, 10 beta maps corresponding to each motion direction, each sensory modality and each hand position were used for Multivariate Pattern Analyses (MVPA).

**Figure 2.**
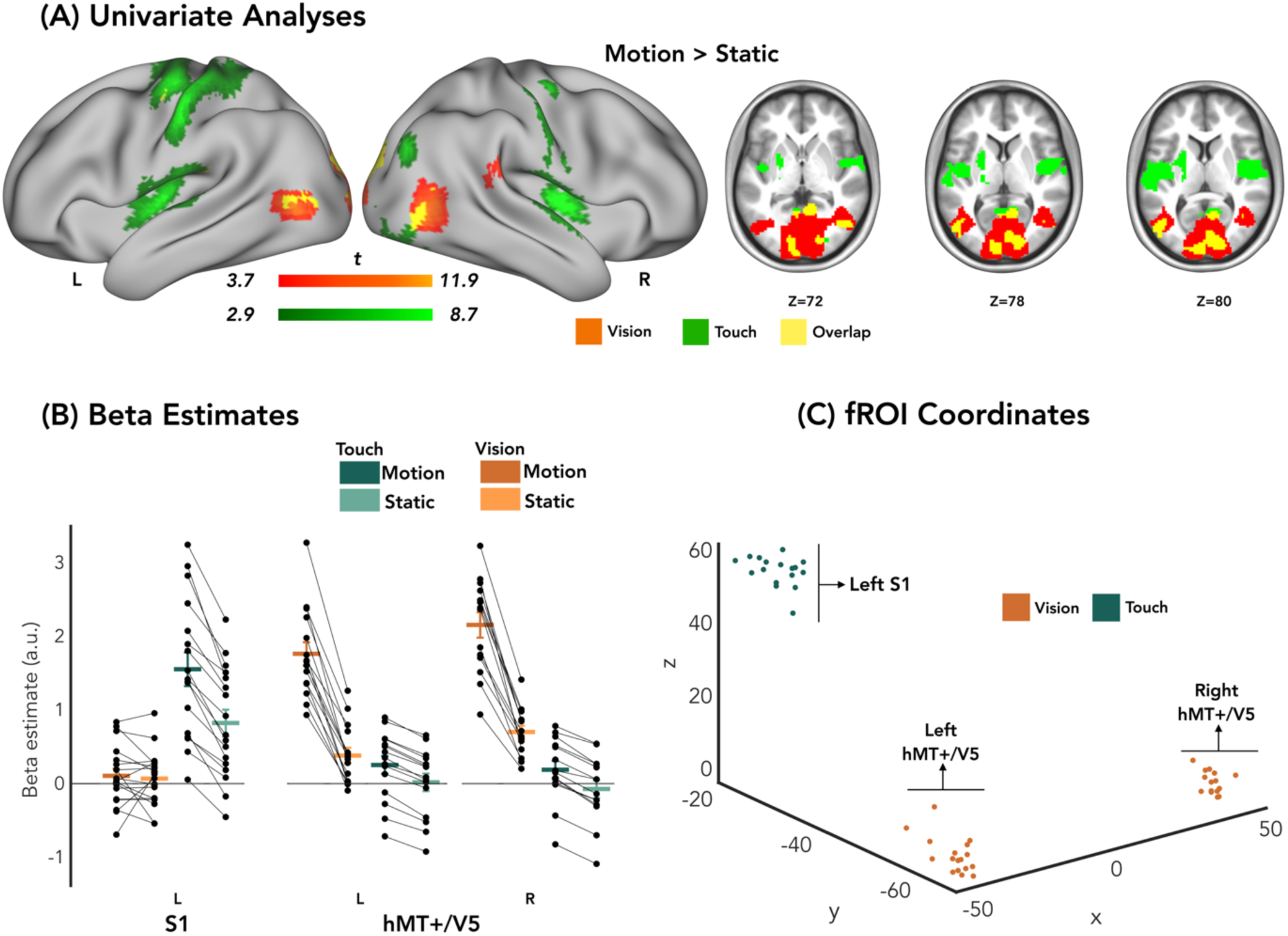
**functional Regions-of-Interest (fROIs) and associated univariate activity.** (A) Group-level univariate analyses of visual (orange) and tactile (green) motion > static contrasts. There is an overlap between the visual and tactile motion selectivity in the hMT+/V5 (yellow). (B) Beta estimates from the visual and tactile motion localizer in each individually-defined fROIs showing visual (orange) and tactile (green) motion selectivity. Tactile beta estimates in hMT+/V5 are from the individually overlapping visual and tactile motion selective response in hMT+/V5. Betas are plotted for illustration purposes only, for statistics please see the appropriate section in the text. Error bars represent the standard error mean (C) MNI coordinates of the individually-defined spherical functional ROIs.

### Stimuli

#### Visual Stimuli

Visual stimuli (Figure 1A) were presented on a black screen with a frame rate of 60 Hz and a resolution of 1920 x 1200 pixels, measuring 70 cm in width and 43.75 cm in height. Participants viewed the screen from a distance of 170 cm through a mirror mounted on the MRI head coil. The stimuli consisted of 300 white random dot kinematograms (RDKs) displayed within an invisible circular aperture, which was centered around a white fixation cross and occupied the full height of the screen (15° of visual angle). The dots could either remain static or move in one of four translational directions: left, right, up, or down, as well as radially receding or approaching. Each dot measured 0.25° in size, had a limited lifetime of 200 ms (Bex et al., 2003), and moved at a speed of 15°/s. The origin of each dot was randomized on each trial.

#### Tactile Stimuli

Tactile stimuli (Figure 1A) were delivered manually using a custom-built, 30 cm wide MRI-compatible brush. The trained experimenter (author Iqra Shahzad), who was specifically trained to deliver tactile stimuli with consistent timing and frequency, brushed the right palm of the participants following computer-controlled auditory cues, which were delivered via MRI-compatible earphones (S14 insert earphones, SensiMetrics, USA; http://www.sens.com/). Each swipe lasted approximately 1s. Participants were instructed to keep their eyes closed throughout all tactile runs, and compliance was monitored by the experimenter.

### Experiments

#### Visual Motion Localizer

One run of visual motion localizer (Figure 1B) was implemented to localize hMT+/V5 both at the group-level and in each participant individually (Huk et al., 2002; Rezk et al., 2020). The hMT+/V5 region, located in the medial temporal area responds preferentially to moving stimuli compared to static stimuli. Since the visual stimuli were presented at fixation in the center of the screen, the localizer defined the whole hMT+/V5 complex which includes both middle temporal area (MT) and medial superior temporal area (MST) (see below for a specific MT/MST localizer). Visual motion and static conditions were generated using white random dot kinematograms (RDK) on a black background. Each run lasted 4 minutes and 51 seconds, starting with an initial 5.25 seconds of a blank screen and ending with 14 seconds of blank screen. Within the run, there were 12 blocks of alternating motion and static conditions, with the starting block counterbalanced across participants. The blocks were separated by an inter-block interval (IBI) of 5 to 7 seconds, chosen randomly. Each block lasted 17.4 seconds and included 6 trials of 2.4 seconds each, separated by an inter-stimulus interval (ISI) of 0.5 seconds. In the motion blocks, the dots moved in one of four directions – left, right, up, and down. Within a motion block, a trial consisted of an axis of motion direction – horizontal (leftward movement of 1.2s followed by rightward movement of 1.2s) or vertical (upward movement of 1.2s followed by downward movement of 1.2s) axis. In the static blocks, each trial (2.4s) consisted of the sequential presentation of two static dots configurations, each lasting for 1.2s (following the cycle of motion direction change). Participants were asked to fixate on the central fixation cross. In the motion blocks, they were asked to detect a repeated direction within a trial, for example, leftward followed by a leftward motion stimulus. In the static blocks, they were asked to press when the dots did not change configuration in the two sequential presentations. The number of targets in each block varied between 0 and 2, randomly distributed and balanced across motion and static conditions.

#### MT/MST visual motion localizer

Using visual radial-motion stimulation, a separate localizer was implemented to divide the hMT+/V5 complex into subregions that may be identified as homologs to a pair of macaque motion-responsive visual areas: the middle temporal area (MT) and the medial superior temporal area (MST) (Huk et al., 2002). MT, thought to have smaller retinotopically organized receptive fields should not respond to peripheral ipsilateral stimulation. MST, known to have larger receptive fields, is assumed to respond to peripheral visual motion stimulation in both ipsilateral and contralateral visual hemifields (Huk et al., 2002). One run of MT/MST localizer (∼7min) (Figure 1B) consisted of 20 blocks of stimulation: 10 blocks presented radial motion in the left hemifield, and the following 10 blocks presented the radial motion in the right hemifield, while the fixation was maintained in the opposite side of the screen i.e., right and left respectively. The order of presentation of the motion stimuli in the left or right hemifield was counterbalanced across participants. Within a block, there were 6 trials, where each trial consisted of receding (0.6s) followed by approaching (0.6s) RDKs. The dots were restricted to a circular aperture of radius 7° visual angle, with its edge 10° from fixation. The run started with a blank screen for 5s and ended with a blank screen for 14s. The inter-block interval (IBI) was 10s including trials during which participants had to change the fixation from one side to the other. Participants were asked to fixate their eyes on the fixation cross and detect changes in its color (white to red). The number of targets (range: 0-2 in each block) were randomized and balanced across conditions.

#### Tactile Motion Localizer

A tactile motion localizer (Figure 1B) was implemented to identify regions that respond preferentially to tactile motion. Each participant performed two runs of the tactile motion localizer (10min 49s each). Each run started with a blank screen of 5s and ended with a 14s pause. It included 20 blocks of alternating motion and static conditions with the starting condition counterbalanced across participants. The blocks were separated with an inter-block interval (IBI) chosen randomly between 5-7 s. Each block (19.5s) included 6 trials of 2s each and an inter-stimulus interval (ISI) of 0.5s. A motion trial consisted of a brush swipe on the right palm from pinky-to-thumb (1s) followed by thumb-to-pinky (1s) representing one somatotopic axis of motion, or from finger-to-wrist (1s) followed by wrist-to-finger (1s) representing the other somatotopic axis of motion. In the static trials, the brush was touched on the right palm sequentially across two somatotopic orientations: thumb-pinky (1s) and finger-wrist (1s). Participants kept their eyes closed throughout the run. In the motion blocks, they were asked to detect a repeated direction within a trial, for example, pinky-to-thumb followed by a pinky-to-thumb motion. In the static block, the participants were asked to detect the trials where the brush was touched in the same orientation consecutively. The number of targets within a block varied between 0 and 2 with the position of the target in the blocks randomized and balanced across conditions.

#### Directional Motion Decoding Experiments

The directional motion experiments (Figure 1C) were conducted to characterize the pattern of neural activity underlying perception of visual and tactile motion directions across two hand positions – HU and HD using multivariate pattern analyses (MVPA). These experiments involved the presentation of directional motion stimuli across 30 runs, such that 10 runs were performed for each sensory modality and each hand position - i.e. visual, tactile HU, tactile HD. The order of the 3 types of runs was counterbalanced across participants. Note that the participants were allowed to rest their right hand in a comfortable position during the visual runs and the motion stimuli used in these experiments were identical to those used in the localizer tasks. Each run lasted 3 min 18s, beginning with a 5s blank screen ending with a 14 s blank screen. Within each run, two directional motion stimuli were presented four times each in a randomized order. Thus, there were six motion conditions across 30 runs (visual: horizontal (Vh) and vertical (Vv); tactile: hand up pinky-thumb (HUpt) and finger-wrist (HUfw); tactile: hand down pinky-thumb (HDpt) and finger-wrist (HDfw)). Each block (17s) comprised 7 trials of a directional stimulus along one axis-of-motion (either pinky-thumb or finger-wrist axis for tactile runs; either horizontal or vertical axis for visual runs), with an ISI of 0.5 s and an IBI chosen randomly between 5 and 7s.

#### MRI acquisition

A whole-brain T1-weighted anatomical scan was acquired (3D-MPRAGE; 1.0 x 1.0 mm in-plane resolution; slice thickness 1mm; no gap; inversion time = 900 ms; repetition time (TR) = 2189,12 ms; echo time (TE) = 2.96 ms; flip angle (FA) = 8°; 156 slices; field of view (FOV) = 256 × 256 mm^2^; matrix size= 256 × 256).

Functional data for the localizers was acquired with T2*-weighted scans acquired with Echo Planar and gradient recalled (EP/GR) imaging (2.6 x 2.6 mm in-plane resolution; slice thickness = 2.6mm; no gap; Multiband factor = 2; TR = 1750ms, TE = 30ms, 58 interleaved ascending slices; FA = 75°; FOV = 220 × 220 mm^2^; matrix size= 84 × 84). The number of volumes acquired in the visual motion localizer, tactile motion localizer and MT/MST visual localizer were 167, 372 and 282 volumes respectively.

Functional data for the directional motion decoding experiments were acquired with a T2*- weighted scans acquired with Echo Planar and gradient recalled (EP/GR) imaging (2.3 × 2.3 mm in-plane resolution; slice thickness = 2.3mm; no gap; Multiband factor = 2; TR = 1750ms, TE = 35ms, 52 interleaved ascending slices; FA = 75°; FOV = 216 × 216 mm^2^; matrix size= 94 × 94). A total of 117 volumes were collected for each tactile HU and HD runs, while 114 volumes were collected for each visual runs.

### Behavioral data analyses

The performance of participants in all the different experiments was analyzed and reported as mean accuracy of detecting targets with standard error. Two-sample t-tests were computed to check if there were differences in detection performance between two conditions within an experiment.

### fMRI Preprocessing

The structural and functional data were pre-processed with BidSpm (v2.2.0; https://github.com/cpp-lln-lab/bidspm; Gau et al. 2022) using Statistical Parametric Mapping (SPM12 - 7771; Wellcome Center for Neuroimaging, London, UK; https://www.fil.ion.ucl.ac.uk/spm; RRID:SCR_007037) and MATLAB R2020b on a macOS (macOS Monterey Version 12.6) computer. The preprocessing of the functional images was performed in the following order: slice timing correction with reference to the middle slice; realignment and unwarping; segmentation and skull-stripping; coregistration of functional and anatomic data; normalization to a template in Montreal Neurologic Institute (MNI) space and smoothing with all default options implemented in BidSpm. Slice timing correction was performed taking the 29th (middle) slice as a reference (interpolation: sinc interpolation). Functional scans from each participant were realigned and unwarped using the mean image as a reference (SPM single pass; number of degrees of freedom: 6; cost function: least square) (Friston et al., 1995). The anatomical image was bias field corrected, segmented and normalized to MNI space (target space: IXI549Space; target resolution: 1 mm; interpolation: 4th degree b-spline) using a unified segmentation. The tissue probability maps generated by the segmentation were used to skull-strip the bias corrected image removing any voxel with p (gray matter) + p (white matter) + p (CSF) > 0.75. The mean functional image obtained from realignment was co-registered to the bias corrected anatomical image (number of degrees of freedom: 6; cost function: normalized mutual information) (Friston et al., 1995). The transformation matrix from this coregistration was applied to all the functional images. The deformation field obtained from the segmentation was applied to all the functional images (target space: IXI549Space; target resolution: equal to that used at acquisition; interpolation: 4th degree b-spline). Preprocessed functional images spatially smoothed using a 3D gaussian kernel (FWHM of 6mm for the localizers and 2mm for the directional motion decoding experiments). This section of preprocessing fMRI data was automatically generated using BidSpm (Gau et al., 2022).

### Univariate Analyses

For both visual and tactile motion localizer, a general linear model (GLM) was fitted for every voxel with the motion and the static conditions as regressors of interest in each subject. Each regressor was convolved with SPM canonical hemodynamic response function (HRF). The 6 head motion parameters (3 translation and 3 rotation parameters) were included in the model as regressors of no interest. The contrast (motion > static) was computed for every subject individually to identify functional regions-of-interest (fROIs) responding preferentially to motion stimuli in vision and touch. In addition, these individual contrasts were further smoothed with a 6 mm FWHM kernel and entered a random effects model for a standard second-level group analysis. Group-level statistical significance was assessed either using family-wise error (FWE) correction across the whole brain at a significance level of p < 0.05 or using small volume correction (SVC) within small spherical volumes (10 mm radius) around functional ROI coordinates. (Table S 1 and Table S2). Regions were labelled using the Automated Anatomical Labelling 3 (AAL3) atlas (Rolls et al., 2020) available in SPM12.

A similar approach was adopted for the univariate analyses of MT/MST visual motion localizer. A GLM was fitted for every voxel with left-hemifield motion and right-hemifield motion conditions as regressors of interest in each subject. Each regressor was convolved with SPM canonical hemodynamic response function (HRF). The 6 head motion parameters (3 translation and 3 rotation parameters) were included in the model as regressors of no interest. The contrasts (left-hemifield motion > baseline and right-hemifield motion > baseline) were computed for every subject individually. The conjunction of these contrasts revealed the MST area in each individual.

### Functional Regions of Interest (fROI) definition

The individually defined motion selective functional regions of interest (fROIs) were created following the Group-Constrained Subject-Specific (GSS) method, which algorithmically detects fROIs that are systematically activated across subjects. This method is similar to a random effects analysis, but is more tolerant of anatomical variability across subjects, adding sensitivity and selectivity to the results. The GSS method has been successfully used to define fROIs preferential for various visual categories (Julian et al., 2012), language (Fedorenko et al., 2010) and visual and auditory motion (Battal et al., 2022; Dormal et al., 2016; Dormal & Collignon, 2011; Rezk et al., 2020). The visual motion selective area, hMT+/V5 was identified from the visual motion localizer using the contrast motion > static. First, hMT+/V5 was identified bilaterally at the group-level as the cluster of voxels (p<0.05 FWE) in the middle-temporal gyrus. The group-level spherical fROI mask (radius 12mm) was drawn around the group-level peak coordinate in each participant. Within this group-constrained mask, individually identified peak of motion selectivity (motion > static) was identified and an individual spherical fROI masks was selected (radius 8mm; ∼ 176 voxels).

Further, MT/MST visual motion localizer was used to identify and create fROI masks of the sub-divisions of the hMT+/V5: MT and MST. MST was defined as the part of hMT+/V5 that responds to ipsilateral and contralateral visual motion stimulation. Therefore, individually localized clusters of MST voxels were constrained by the individually localized clusters of hMT+/V5 and used as fROIs masks in further analyses. Individual MT masks were defined as the part of hMT+/V5 that prefers contralateral over ipsilateral motion and therefore excludes MST.

A similar approach was adopted to identify tactile motion selective areas. Group level analyses first revealed the presence of tactile motion selective response in the postcentral gyrus and middle temporal cortex. As in the visual motion localizer, a 12mm spherical fROI mask was drawn around the group-level peak coordinate to constrain individual subject activity. Ultimately, an 8mm-radius spherical fROI mask was drawn around the individually identified peak coordinates. Spherical masks of tactile motion selective areas in the middle temporal cortex were created in the same manner and were referred to as MTt (Middle Temporal tactile).

To foreshadow the univariate results, tactile motion selective areas overlapped with the visual motion selective hMT+/V5 in the middle temporal cortex, showing that a part of hMT+/V5 is engaged in tactile motion processing. Since the individually defined hMT+/V5 and MTt overlapped with each other (see Table S 4 for dice coefficients), we reported hMT+/V5 results in the main findings while presenting MTt results as supplement (Figure S 2). The analyses in hMT+/V5 and MTt produced similar results (Figure 3 and Figure S 2), which was expected given the substantial overlap between these regions (Table S 4).

**Figure 3.**
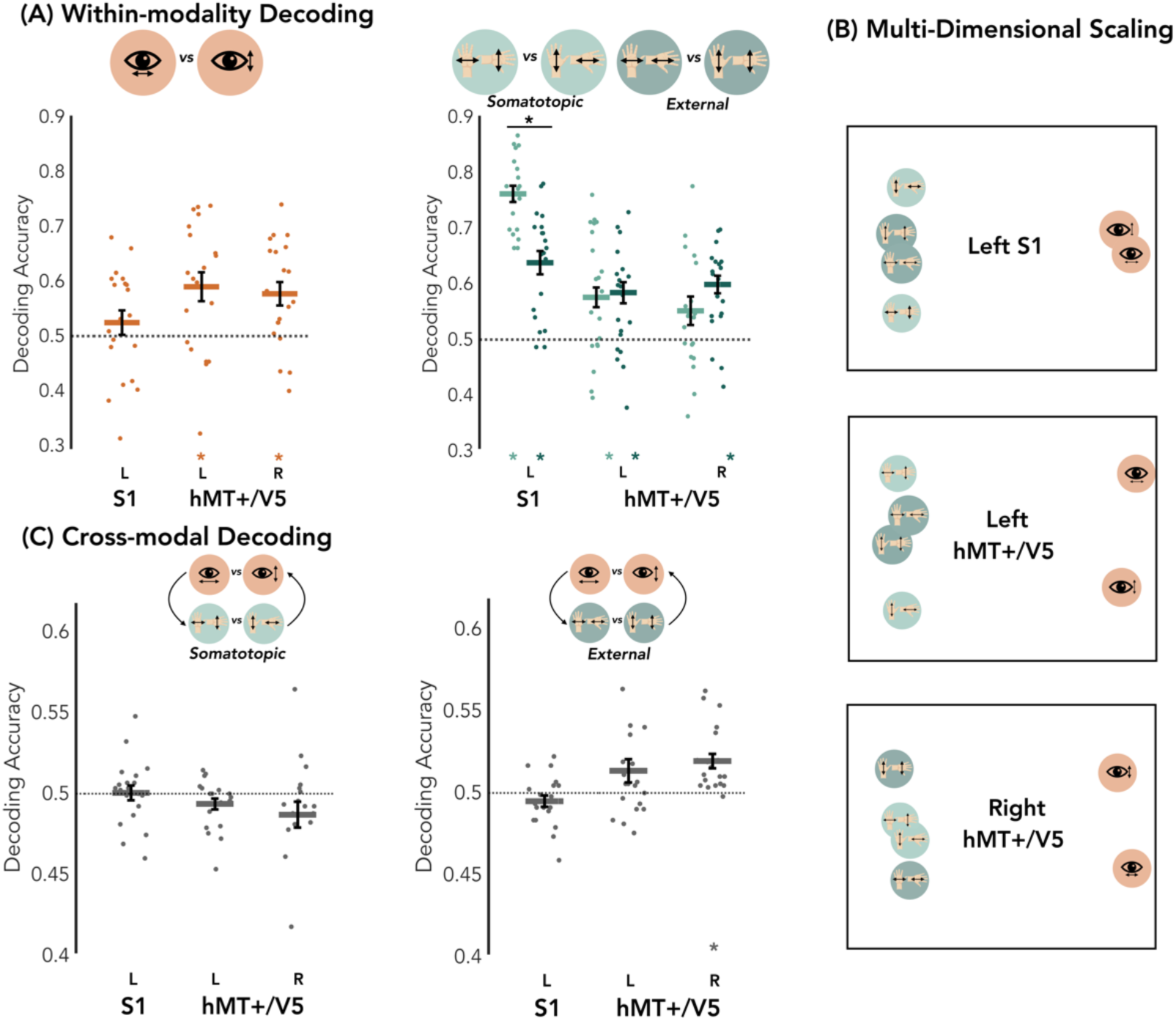
**fROI-based MultiVariate Pattern Analyses (MVPA).** (A) Within-Modality Decoding. Binary classification of visual (left panel) and tactile (right panel) motion directions across the two frames of references in each fROI. The y-axis represents the decoding accuracy with chance level at 0.5. Results are FDR corrected (*p<0.05) for the number of fROIs. (B) Multi-Dimensional Scaling (MDS). Visualisation of the relative distance between the patterns of the visual and tactile motion directions across the two frames of reference in each fROI. (C) Cross-Modal Decoding. Cross-modal classification between visual and tactile motion directions across the two frames of reference – somatotopic (left panel) and external (right panel) within each fROI. The classifier is trained on visual motion-directions and tested on tactile motion-directions, and vice-versa, with the decoding accuracies subsequently averaged (see supplemental material for each direction separated). Results are FDR corrected (*p<0.05) for the number of ROIs.

### Multivariate Pattern Analyses (MVPA)

Multivariate pattern analyses (MVPA) were carried out to investigate the representation of visual and tactile motion-directions across different hand postures in each subject. They were performed on the GLM output of directional motion decoding experiments. A GLM was fitted for each run and in each subject. Depending on the type of run, the regressors of interest were the two axis-of-motion stimuli (visual: horizontal (Vh) and vertical (Vv); tactile: hand up pinky-thumb (HUpt) and finger-wrist (HUfw); tactile: hand down pinky-thumb (HDpt) and finger-wrist (HDfw)) and the 6 head motion parameters were included as regressors of no interest. One beta map corresponding to each condition from each run was obtained. In total, we computed a total of 60 beta maps for each subject (10 for Vh, Vv, HUpt, HUfw, HDpt, HDfw).

To improve the SNR of each beta map used for decoding, the 10 beta maps within each experimental condition (eg. Vh) were randomly paired and averaged to generate 5 pseudorun-beta maps. Henceforth, the general procedure involved binary classification of the different experimental conditions using linear Support Vector Machine (SVM) (C parameter = 1) implemented in CoSMoMVPA (http://www.cosmomvpa.org/) (Oosterhof et al., 2016) and LIBSVM (http://www.csie.ntu.edu.tw/;cjlin/libsvm) (Chang & Lin, 2011) following a leave-one-run-out cross-validation scheme. Since there were 45 unique combinations of the pseudoruns, this procedure was repeated 45 times.

The binary decoding of different experimental conditions was performed within the fROI created from the localizer experiments. An ANOVA-based feature selection method was used to keep the most informative 100 features/voxels within a fROI. This feature selection process not only ensures a similar number of voxels within a given region across participants, but, more importantly, identifies and selects voxels that are carrying the most relevant information across categories of stimuli (Cox & Savoy, 2003; De Martino et al., 2008); therefore, minimizing the chance to include voxels that carry noises unrelated to our categories of stimuli. The classifier was trained on 4 pseudorun-betas and tested on 1 pseudorun-beta. This step was repeated 5 times (N-fold cross-validation) where in each fold the classifier was tested on a different run. The performance of the classifier was reported as the classification/decoding accuracy for each subject by averaging the decoding accuracies across all cross-validation folds. Finally, the mean decoding accuracy was reported across all the subjects with its standard error. Above chance (0.5 for binary classification) decoding accuracy would indicate presence of axes of motion information.

Statistical significance of the mean decoding accuracies within a fROI was assessed using non-parametric methods that combine permutations and sign flipping (Nichols et al., 2005; Winkler et al., 2016; Xu et al., 2023). First, the observed mean decoding accuracy was calculated across the subjects. Then, 10,000 iterations of a sign-flipping procedure were performed to generate a group-level null distribution. In each iteration, the decoding accuracy of each subject was randomly assigned a reverse sign, and the mean decoding accuracy was recalculated for the group. This gave us 10,000 mean decoding accuracies which formed the group-level null distribution. Statistical significance was then determined by comparing the observed mean decoding accuracy to the null distribution, with the p-value defined as the proportion of null distribution values greater than or equal to the observed mean. To account for multiple comparisons across fROIs, all *p* values were corrected using false discovery rate (FDR) correction (Benjamini & Hochberg, 1995).

### Within-modality decoding

To test the presence of visual motion direction information in motion-selective areas, binary classification of visual horizontal and visual vertical directions was implemented in the fROIs defined from the localizers– left somatosensory cortex, left and right hMT+/V5. Since hMT+/V5 (defined from the visual motion localizer) and MTt (defined from the tactile motion localizer) overlapped with each other, the decoding of visual motion information in MTt is presented in supplemental material. To further investigate the visual motion selectivity in the sub-divisions of hMT+/V5, classification was performed in the MST and MT fROIs masks separately.

Since tactile motion directions can be coded either in a somatotopic or in an external frame of reference depending on the hand positions, the four experimental conditions were rearranged to perform two different classifications in the motion-selective fROIs. The somatotopic frame of reference was decoded by classifying pinky-thumb direction versus finger-wrist direction pooled across the hand position. The external frame of reference was decoded by classifying horizontal (pinky-thumb in HU position pooled with finger-wrist in HD position) versus vertical (finger-wrist in HU position pooled with pinky-thumb in HD position) directions. To further study the recruitment of tactile frame of references in the different ROIs, the decoding accuracies were entered into a Linear Mixed Model (LMM) using the *lmer* function implemented in R (R Core Team, 2023). The frame of references (somatotopic, external) and the fROIs (left S1, left hMT+/V5, right hMT+/V5) were entered as fixed effects and subjects as a random effect. Finally, the tactile motion representation was investigated in the same manner in the MST, MT and the MTt fROIs separately.

### Multi-Dimensional Scaling

Multi-Dimensional Scaling (MDS) was used to visualize the underlying spatial structure of the different motion directions across the sensory modalities in each fROI. It allowed us to visualize the relative distance between the different pairs of motion directions across the sensory modalities. To do so, representational similarity analysis (RSA) (Kriegeskorte, 2008; Kriegeskorte et al., 2008) was first used to assess the underlying dissimilarities across the 6 different experimental conditions (visual horizontal, visual vertical, tactile pinky-thumb in somatotopic frame of reference, tactile finger-wrist in somatotopic frame of reference, tactile horizontal in external frame of reference, tactile vertical in external frame of reference). The dissimilarity matrix was created using the pairwise distances (metric “correlation distance” in CoSMoMvpa) for each subject and then averaged across subjects for a single fROI. The averaged dissimilarity matrices were visualized on an abstract cartesian map of 2-dimensions using DISTASIS (Abdi et al., 2005).

### Cross-modal Decoding

To test the presence of shared axes of motion information across the senses, we performed crossmodal binary decoding of visual and tactile motion axes of motion within each fROIs – left somatosensory cortex, left hMT+/V5 and right hMT+/V5 and separately for MT, MST and MTt. Here, the classification was implemented using a similar cross-validation scheme as described above, except that the training set was chosen from one modality and the testing set from another modality. Also, note that prior to the cross-modal classification, the patterns of each motion direction corresponding to each modality were demeaned to equate their mean activity. Since the tactile motion directions could be decoded in two different frames of reference, crossmodal decoding was performed between (a) visual motion directions and tactile motion directions in a somatotopic frame of reference, and (b) visual motion directions and tactile motion directions in an external frame of reference. Note that the analytical procedure was the same for both (a) and (b) except the way the tactile stimulation was assigned to different classes. Finally, statistical inference of the mean decoding accuracies across both the direction of training and testing sets (train on vision, test on touch and vice versa) within each fROI are reported. The crossmodal decoding for the different directions of train and test sets is reported in supplemental material (Figure S 1).

### Searchlight Analyses

In addition to the fROI-based MVPA, we implemented whole-brain searchlight decoding analyses (Kriegeskorte et al., 2006), following the approach used in (Rezk et al., 2020) to identify regions representing visual and tactile motion information outside of fROIs defined using the visual and tactile motion localizers. A sphere (3-voxel radius; ∼113 voxels) was moved across the brain volume, where in each step, each voxel became the center of the searchlight sphere. The decoding accuracy computed within the sphere was assigned to the central voxel, generating a whole-brain map of decoding accuracies for each participant. Individual accuracy maps were smoothed using an 8 mm FWHM gaussian kernel and entered in a second-level group analysis. Group-level maps were corrected using small-volume correction (SVC) within 10-mm radius spheres centered on hMT+/V5 locations, which were independently identified using the visual motion localizer (left hMT+/V5: −47, −70, 5 mm; right hMT+/V5: 47, −65, 3 mm). Since group-level univariate analyses did not reveal parietal areas as motion selective, we applied SVC at the coordinates of parietal regions previously shown to be selective for tactile motion (Summers et al., 2009) (MNI coordinates left parietal area: - 30, −53, 49 mm; right parietal area: 41, −55, 54 mm).

## Results

### Behavioral Responses

We assessed the behavioral performance of each participant in all the experiments, measuring their ability to detect targets in each condition. Table S 1 presents the mean accuracy in each task. Participants performed consistently well across all conditions, with no significant differences between motion and static conditions in the localizer experiments or between conditions (Vh and Vv; HUpt and HUfw; HDpt and HDfw) in the main decoding experiments (Wilcoxon signed rank test; all *p* > 0.05). This lack of difference in behavioral performance between conditions suggests that the observed motion selectivity in the localizers and the decoding results in the directional motion decoding experiments cannot be attributed to differences in performance or attention.

### Visual Motion Localizer

Table S 1 lists the regions showing preferential responses to visual motion at a threshold *p* = 0.05 FWE corrected which are illustrated on Figure 2A. We found clusters of voxels in the left and right middle temporal gyrus (hMT+/V5) extending bilaterally to lingual gyrus (V1) and left V3a. The hMT+/V5 clusters were observed at the intersection of the ascending limb of inferior temporal sulcus and lateral occipital sulcus (Dumoulin, 2000). We did not observe regions such as VIP and LIP in the parietal cortex to be visual motion selective using our localizer approach which is also consistent with previous studies (Battal et al., 2022; Dormal et al., 2016; Rezk et al., 2020; Zeki et al., 1991).

### Tactile Motion Localizer

We adopted a similar approach to identify tactile motion selective areas from the whole brain univariate analyses and computation of the contrast tactile motion > tactile static (Figure 2A, Table S 2). Consistent with previous studies (Ricciardi et al., 2007; Summers et al., 2009; van Kemenade et al., 2014), we observed preferential response to tactile motion in the left post central gyrus (left primary somatosensory cortex), bilateral rolandic operculum (secondary somatosensory cortex) and in the middle temporal gyrus. An overlay of the group-level contrasts (motion > static) revealed an overlap between the visual and tactile motion selective regions (Figure 2A). More precisely, 32.21% of voxels of visually defined left hMT+/V5 and 42.55% of voxels in right hMT+/V5 showed tactile motion selectivity. Similar to the visual motion localizer, we did not observe regions in the parietal cortex to be tactile motion selective.

To further investigate tactile motion selectivity within the individually localized bilateral hMT+/V5, we compared the beta parameters for tactile motion and static conditions within each individual’s hMT+/V5 fROI (Figure 2B). One-tailed paired t-tests revealed that responses to tactile motion conditions were significantly higher than the static conditions in both fROIs (left hMT+/V5 (t (17) = 1.99, p = 0.03, d = 0.24) and right hMT+/V5 (t (17) = 2.15, p = 0.02, d = 0.17). The fROI left primary somatosensory cortex being tactile motion selective did not show visual motion selectivity (t (17) = 0.47, p = 0.32, d = 0.09).

### MT/MST Localizer

We identified subregions of hMT+/V5, which are presumed homologs to the macaque middle temporal (MT) area and the medial superior temporal (MST) area, in each individual (Huk et al., 2002). In each subject, we computed the conjunction (AND) (Nichols et al., 2005) of the contrasts left-hemifield motion > 0 and right-hemifield motion > 0 at a threshold of *p* = 0.001 uncorrected or *p* = 0.05 FWE corrected, and constrained the activity to the individually localized clusters of hMT+/V5 (from the visual motion localizer) to identify MST area. Table S 3 shows the number of voxels in the identified clusters. Following this, MT voxels were defined as part of individually localized clusters of hMT+/V5 which do not belong to MST. We were able to reliably localize these regions in 14 out of the 18 tested participants. Figure 5A shows the identified clusters of MST and MT in one exemplar subject.

**Figure 4.**
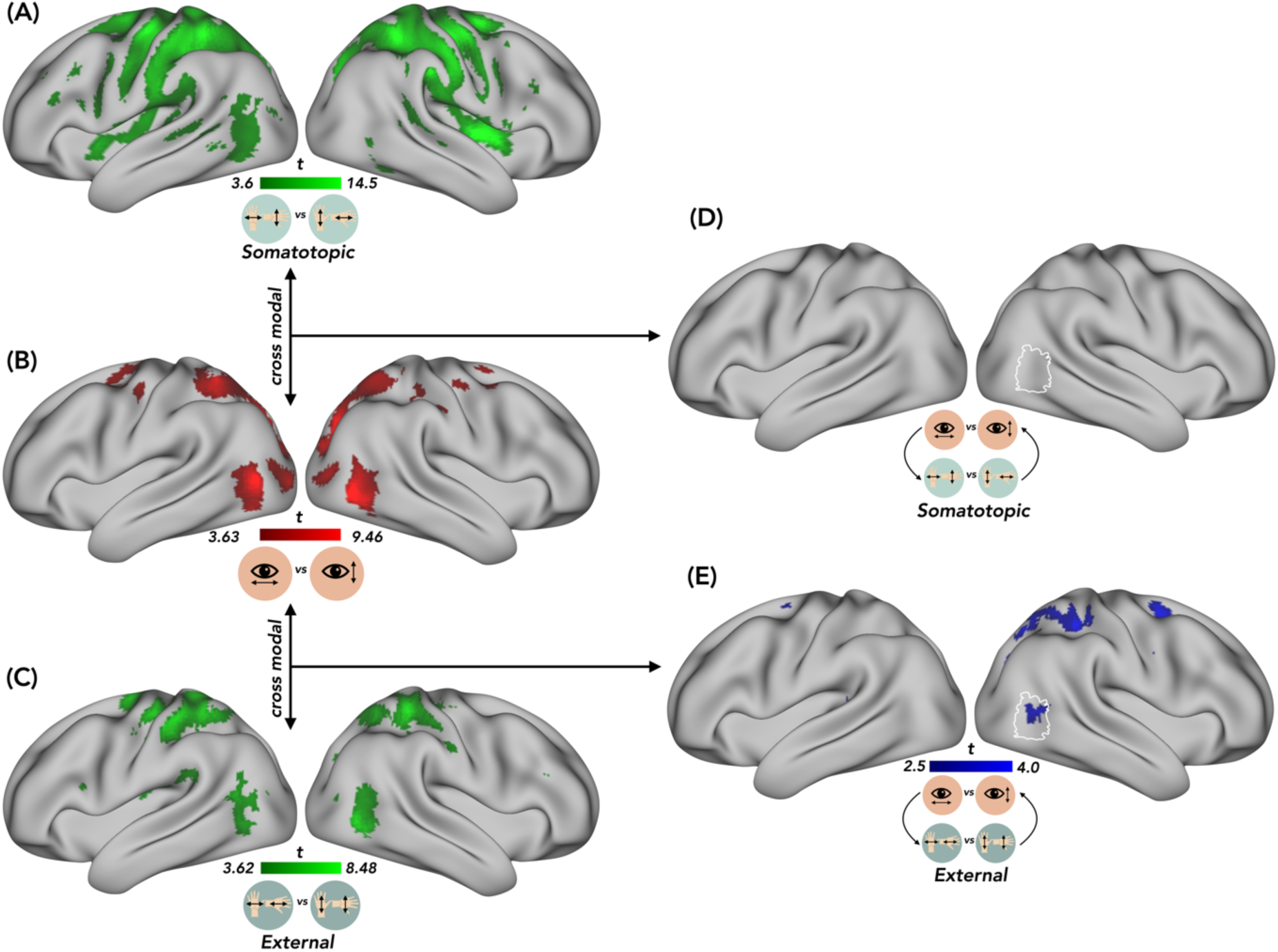
**Searchlight Analyses.** (A) Within-modality decoding of tactile motion directions defined in the somatotopic frame of reference. (B) Within-modality decoding of visual motion directions. (C) Within-modality decoding of tactile axis-of-motion directions defined in the external frame of reference. (D) Cross modal decoding between visual and tactile motion directions defined in the somatotopic frame of reference. (E) Cross modal decoding between visual and tactile motion directions defined in the external frame of reference.

**Figure 5.**
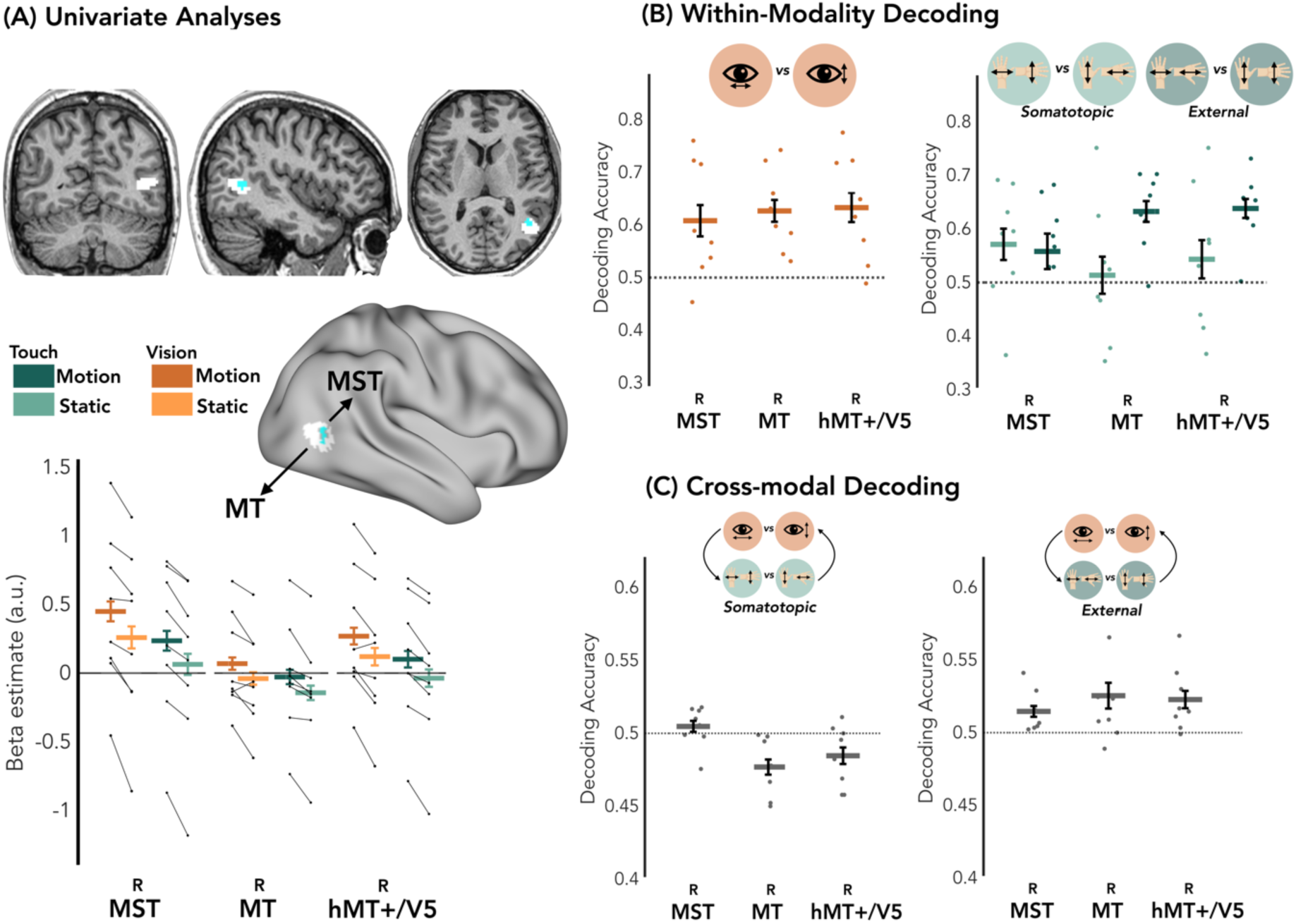
**Analyses in the MT/MST.** (A) Univariate Analyses. Example of an individually localized MST and MT from the MT/MST visual motion localizer (top). Beta estimates of visual – motion and static, and tactile – motion and static in the individually localized right MST, right MT and right hMT+/V5 (bottom). (B) Within-modality decoding of visual and tactile motion directions across the two frames of reference in the right MST, right MT and right hMT+/V5. (C) Cross-modal decoding between visual and tactile motion directions across the two frames of reference – somatotopic (left panel) and external (right panel) within each fROI. In the left panel, the tactile motion directions are defined in somatotopic frame of reference and in the right panel, the tactile motion directions are defined in the external frame of reference.

To investigate tactile motion selectivity within the individually localized MST and MT, we extracted tactile motion and static mean beta parameters from the univariate analyses of tactile motion localizer and compared them within each individual’s MST and MT mask (Figure 5A). We performed one-tailed paired t-tests (*p*-values subjected to Bonferroni correction for multiple comparisons) to compare the motion and static conditions within each fROI. We obtained significantly higher beta parameters for motion conditions as compared to the static conditions in all the fROIs (left MST (t(13) = 3.37, *p* = 0.005, *d* = 0.39), left MT (t(13) = 3.13, *p* = 0.005, *d* = 0.18), right MST (t(13) = 4.76, *p* < 0.001, *d* = 0.37), right MT (t(13) = 3.49, *p* = 0.005, *d* = 0.25)).

### fROI-based MVPA

#### Functional Regions of Interest (fROIs)

We performed whole-brain univariate analyses for independent visual and tactile motion localizers (contrast motion>static) to acquire the group-level coordinates of bilateral hMT+/V5 and left S1, selective to visual and tactile motion, respectively. We first used the MNI group coordinates: left S1 [−44, −2, 60 mm]; left hMT+/V5: [−42, −65, 4 mm]; right hMT+/V5: [48, −63, 1 mm] to create group-level spherical fROI masks of radius 12mm. Following this, we identified individual peak coordinates, constrained by the group-level fROI masks, to create spherical fROI masks of radius 8mm. Table S 5 shows the MNI coordinates of the individual spherical fROI.

Since we observed preferential response to tactile motion from the tactile motion localizer in the hMT+/V5 region (for overlap, see Figure 2A), we performed separate MVPA in hMT+/V5 (defined using visual motion localizer) and MTt (defined using tactile motion localizer). Since the individually defined hMT+/V5 and MTt overlapped with each other (for dice coefficients see Table S 7), results in hMT+/V5 are presented in the main manuscript and in MTt in supplemental; both fROIs yield similar results.

Foreshadowing our MVPA results, we observed significant crossmodal decoding only in the right hMT+/V5 (Figure 3C). Therefore, we performed MVPA in the individually identified clusters of MST and MT to further investigate the visual and tactile motion representations across different frames of reference solely in the right hemisphere. However, due to the variability in the size of the fROIs masks across participants (as shown in Table S 1), we chose a subset of 8 subjects which had at least 30 voxels for MVPA. This allowed us to keep the same number of features/voxels for MVPA across regions and subjects.

#### Within-modality decoding

When decoding visual motion axes of motion (vertical versus horizontal), we obtained above-chance (0.5) decoding accuracies in both left (*p* = 0.046 FDR-corrected) and right hMT+/V5 (*p* = 0.046; FDR-corrected), but not in the left S1 (*p* = 0.145; FDR-corrected) (Figure 3A). A separate analysis in MTt (Figure 5A) also revealed significant decoding of visual motion direction (left MTt: *p* = 0.031; FDR-corrected and right MTt: *p* = 0.031; FDR-corrected). In addition, MT and MST (Figure 5B) also suggest the presence of visual motion direction information.

To determine whether the tactile motion directions are represented within a somatotopic frame of reference or an external frame of reference, we performed decoding of tactile motion directions pooled across the hand postures. For somatotopic frame of reference, we classified pinky-thumb vs finger-wrist tactile directions across hand postures. For external frame of reference, we classified horizontal vs vertical tactile directions regardless of the hand postures (see Figure 1A for the design). Decoding accuracies obtained for each frame of reference (somatotopic, external) within the 3 fROIs (left S1, left hMT+/V5, right hMT+/V5) were tested against the chance level (0.5).

The decoding of tactile motion directions in somatotopic frame of reference was significantly above chance in the left S1 (*p <* 0.001; FDR-corrected) and left hMT+/V5 (*p* = 0.024; FDR-corrected). The decoding of tactile motion directions in external frame of reference was above chance in all the three fROIs - left S1 (*p* = 0.009; FDR-corrected), left hMT+/V5 (*p* = 0.013; FDR-corrected), right hMT+/V5 (*p* = 0.007; FDR-corrected) (Figure 3A).

To further assess the recruitment of tactile frames of references within the fROIs, the decoding accuracies were entered into a Linear Mixed Model (LMM). The two frames of reference (somatotopic, external) and the three fROIs (left S1, left hMT+/V5, right hMT+/V5) were entered as fixed effects and participants as random effect. We observed a significant main effect of the fROIs (F (2,95) = 28.01, *p* < 0.001, μ^2^ = 0.35), showing that the primary somatosensory area showed highest decoding accuracy owing to its primary function of processing tactile inputs. There was no significant main effect of the frames of reference (F (1,95) = 2.16, *p* = 0.144, μ^2^ = 0.01). Crucially, there was a significant interaction of fROIs and frame of reference emerged (F (2,95) = 11.25, *p* < 0.001, μ^2^ = 0.17) suggesting that the influence of reference frame on decoding accuracies varied across the fROIs. Post hoc paired t-tests (FDR-corrected) were performed to compare decoding accuracies across the two frames of reference within each fROI. Decoding accuracies were significantly higher in the somatotopic frame of reference than the external frame of reference only in the left S1 (t (19) = 4.21, *p* = 0.0014, *d* = 1.13) supporting a dominant somatotopic representation of tactile motion directions in the primary somatosensory cortex. While the decoding accuracy in the right hMT+/V5 appear descriptively higher for external frame of reference, the difference was not statistically significant (t (19) = −1.37, *p* = 0.278, *d* = −0.43). Similarly, no significant difference was observed in left hMT+/V5 (t (19) = −0.28, *p* = 0.78, *d* = −0.08).

Given the overlap between hMT+/V5 and MTt, we also examined decoding in MTt and observed similar results as that in hMT+/V5. The decoding of somatotopic frame of reference reliably above chance in left MTt (*p* = 0.029) and the decoding of external frame of reference was above chance in both, left MTt (*p* = 0.031) and right MTt (*p* = 0.015) (Figure S 2). Paired t-tests, however, did not reveal significant differences between somatotopic and external coding of tactile motion direction in MTt (*p <* 0.05). Descriptively, the decoding accuracy in right MTt, like right hMT/V5, appeared higher for external than the somatotopic frame of reference.

Finally, we performed decoding of tactile motion directions in different reference frames separately within MST and MT fROIs. Consistent with the results in right hMT+/V5, the individually defined right MT also showed a trend towards higher decoding accuracy of tactile motion direction in external, compared to the somatotopic frame of reference (Figure 5B).

### Multidimensional Scaling

To further investigate the representational structure in left S1 and right hMT+/V5, we used Multidimensional Scaling (MDS) to visualize the relative distances between motion direction conditions in a two-dimensional abstract space (Figure 3B). First, MDS revealed a clear separation between visual and tactile modalities in both fROIs. Second, we observed an interesting organization within the tactile modality. In the left S1, the somatotopically defined directions were more widely spaced than the externally defined directions supporting the trend in decoding accuracies across the tactile frames of reference. In contrast, in right hMT+/V5, externally defined directions were more widely spaced than the somatotopically defined directions, aligning with the trend of higher decoding accuracy for the external frame of reference.

### Cross-Modal Decoding

To further investigate whether the motion information is shared across sensory modalities within our fROIs, we performed crossmodal decoding between visual and tactile motion directions, with the tactile stimuli defined in either (a) somatotopic or (b) external frame of reference (Figure 3C and Figure S 1). When tactile motion directions were defined somatotopically, the crossmodal decoding did not yield significant results in any of the fROIs (*p* > 0.05). In contrast, when tactile motion directions were defined using an external frame of reference, significant crossmodal decoding was observed in the right hMT+/V5 (*p* = 0.013, FDR corrected). This suggests that the motion information is shared across vision and touch in right hMT+/V5, only when the tactile motion directions are represented in an external frame of reference.

Similar results were also found in right MTt where significant crossmodal decoding was obtained only when tactile motion was defined in an external frame of reference (p = 0.032) (Figure S 2).

Interestingly, both right MST and right MT also exhibited the same trend: crossmodal decoding accuracies appeared to increase above chance when the tactile motion directions were expressed in an external reference frame (Figure 5C).

### Searchlight Decoding Analyses

To explore the broader network of regions sensitive to motion-directions across sensory modalities and frames of reference beyond our fROIs, we implemented whole-brain searchlight decoding analyses. Since our visual and tactile motion localizers did not reveal motion selective activity in parietal and frontal areas (Figure1A), the searchlight method allowed us to also further investigate whether these regions play a role in motion direction processing and using which frame of reference.

We performed both within-modal and cross-modal decoding and tested the statistical significance using Small Volume Correction (SVC) within a 10 mm radius sphere. SVC (Table 3) was performed at hMT+/V5 identified independently from the visual motion localizer (left hMT+/V5: −47, −70, 5 mm; right hMT+/V5: 47, −65, 3 mm) and the parietal regions observed from a previous study (Summers et al., 2009) assessing tactile motion selectivity (Talairach coordinates left Intraparietal Area: −30,-50,45 and right Intraparietal Area: 42, −51, 49 were converted to MNI coordinates left Intraparietal Area: −30,-53,49 mm and right Intraparietal Area: 41, −55, 54 mm, using Bioimage Suite Web : (www.bioimagesuite.org) (Lacadie et al., 2008)). We also tested frontal regions implicated in the external representation of touch (Crollen, Lazzouni, et al., 2017; Lloyd et al., 2003) in searchlight crossmodal decoding (left premotor cortex: −46, 8, 46 mm; right premotor cortex: 24, 4, 58 mm).

Group-level searchlight decoding of visual motion directions (horizontal versus vertical; Figure 4B) revealed bilateral clusters in hMT+/V5 (peak coordinate at left: −41.6, −72.9, 5.9 mm and right 43.5, −68.3, −1 mm) and a wider network of clusters in the primary visual cortex (V1) (left Cuneus: −14, −75, 29 mm; right Cuneus: 16, −82, 27 mm) extending through the superior occipital gyrus to superior parietal lobule (left superior parietal gyrus : −28, −50, 50 ; right superior parietal gyrus: 27, −55, 59 mm) and some additional clusters in the frontal lobe (left middle frontal gyrus: −28, −4, 52 mm; right middle frontal gyrus: 34, −2, 52 mm).

We decoded tactile motion directions represented in both somatotopic (Figure 4A) and external frame (Figure 4C) of reference. For the somatotopic frame of reference (Figure 4A) we observed clusters in the left (peak coordinate: −44, −71, 8.2 mm) and right middle temporal gyrus (peak coordinate: 46, −52, −3.3 mm), as well as an extensive network of the primary (left postcentral gyrus) and secondary somatosensory (Rolandic Operculum and Insula) areas. We also observed parietal (superior parietal gyrus), motor (precentral gyrus; supplementary motor area) and frontal (middle superior frontal gyrus; superior frontal gyrus) areas. Decoding of tactile external frame of reference (Figure 4C) revealed clusters bilaterally in the middle temporal area (middle temporal gyrus left: −48, −61, 15 mm and right: 44, −66, 1 mm) and a network of parietal areas (left superior parietal gyrus), primary (left post central gyrus) and secondary (left rolandic operculum, left supramarginal gyrus) somatosensory areas, motor areas (left supplementary motor area, left precentral gyrus) and frontal areas (medial and superior frontal gyrus).

Finally, we performed crossmodal searchlight decoding between visual and tactile motion directions in both frames of reference. No significant clusters were obtained when tactile motion directions were represented in somatotopic frame of reference (Figure 4D). In contrast, when tactile motion directions were defined in the external reference frame (Figure 4E) we observed a distinct network involving the right middle temporal gyrus (48, −59, 8 mm), right superior frontal gyrus (25, −2, 52 mm) and right superior parietal gyrus (18, −66, 52 mm). The cluster in the middle temporal area closely overlapped with the right hMT+/V5 identified from the visual motion localizer (dice coefficient = 0.9091). The cluster in the frontal region encompassed the frontal eye field (FEF) area (Vernet et al., 2014) and the premotor cortex (Lloyd et al., 2003). The cluster in the posterior parietal cortex includes LIPv, LIPd, MIP and VIP based on the Glasser atlas (Glasser et al., 2016).

## Discussion

We investigated where in the brain visual and tactile motion direction representations align across changes in hand posture. Using univariate analysis, we first confirmed that hMT+/V5 - a region traditionally regarded as exclusively visual (Tootell et al., 1995; Zeki et al., 1991) - also responds preferentially to tactile motion directions compared to static stimuli (Blake et al., 2004; Hagen et al., 2002; Ricciardi et al., 2007). We then showed that, similar to vision, directional tactile motion is encoded within hMT+/V5. Critically, by manipulating hand posture, we found that the right hMT+/V5 encodes tactile motion directions in an external frame of reference, aligning representations across both vision and touch. Notably, this cross-modal alignment emerged only when tactile motion directions were remapped into external spatial coordinates. Finally, our results reveal that hMT+/V5 operates within a broader fronto-parietal network that supports the alignment of multisensory motion representations.

Using independent visual and tactile motion localizers, we identified regions that preferentially respond to visual and tactile motion directions at the group (Figure 2A) and individual level (Figure 2C). A significant part of hMT+/V5 exhibited preferential response to both visual and tactile motion directions (43% of right hMT+/V5 voxels), whereas the somatosensory regions responded exclusively to tactile motion. Previous studies have debated whether such motion selective responses to non-visual input were limited to MST but not MT (Beauchamp et al., 2007), or suggested that previous observation of tactile motion signal in hMT+/V5 was due to group averaging (Jiang et al., 2015). Our findings provide unequivocal evidence that parts of hMT+/V5 (Figure 2B), including both MT and MST subdivisions (Figure 5A), preferentially responds to both visual and tactile motion over static stimuli. These preferential activations for tactile motion in regions classically considered purely visual were not a byproduct of spurious group averaging as the tactile motion activity was extracted from individually defined fROIs using independent localizers.

Using MVPA, we investigated the representations of axes of visual and tactile motion within our fROIs and across the different hand postures (Figure 3A). As expected, we found that the visual axes of motion directions can be decoded in bilateral hMT+/V5. In primates, a key organizational feature of this region is that the cortical columns tuned to a specific motion direction are adjacent to those tuned to its 180-degree opposite, implementing an axis-of-motion hyper-columnar organization (Albright, 1984). Such organization likely underlies the ability to decode axes of motion from the neural activity recorded with fMRI in humans (Bartels et al., 2008; Kamitani & Tong, 2006; Schneider et al., 2019; Zimmermann et al., 2011). We further investigated whether tactile motion directions can be decoded from activity patterns in hMT+/V5, and if so, whether they are represented in a somatotopic and/or an external frame of reference. Representing tactile motion in external coordinates requires transforming skin-based inputs by taking hand posture into account (Badde & Heed, 2016; Heed et al., 2015, 2016). This transformation implies that tactile motion directions are abstracted from their original skin location and encoded relative to external space, independent of the body surface on which they were initially sensed. In our study, we did not determine the specific anchor of the external reference frame: while changes in hand posture modified the perceived axis of tactile motion, the positions of potential anchors such as the eyes, head, and trunk remained constant. Therefore, throughout this work, we use the term *external frame of reference* to refer broadly to representations that are non-somatotopic.

The left S1 exhibited significant decoding of tactile motion directions in both somatotopic and external frames of reference, although the representation was stronger in the somatotopic reference frame. This dominant somatotopic encoding aligns with S1’s primary role in processing tactile inputs. Consistent with this, previous studies in macaques have shown that neurons in areas 1, 2, and 3b are highly tuned to specific tactile motion directions (Pei et al., 2011; Pei & Bensmaia, 2014). Recent findings also suggest that area 3b integrates cutaneous and proprioceptive information from the hand (Trzcinski et al., 2023), which may explain the presence of some external reference frame coding in S1, albeit to a lesser extent than somatotopic encoding. The left hMT+/V5 showed significant representation of tactile motion directions in both the reference frames, with no significant differences between them. In contrast, the right hMT+/V5 selectively exhibited a significant representation in the external frame of reference, with no evidence of somatotopic coding. Multidimensional scaling results (Figure 3B) further illustrate these patterns: tactile motion directions defined in external space were more distinctly separated in right hMT+/V5, whereas in left S1 (and to a lesser extent in left hMT+/V5), somatotopically defined motion directions were more distinctly represented. Together, these results indicate that the right hMT+/V5 preferentially encodes tactile motion directions in a non-somatotopic, external frame of reference.

Importantly, we found significant crossmodal decoding between visual and tactile motion directions in the right hMT+/V5 when tactile motion was expressed in an external frame of reference (Figure 3C). This indicates that visual and tactile spatial maps in hMT+/V5 are aligned within a partially shared external reference frame, following the integration of somatotopic coordinates with hand posture information. Our results suggest that spatial selectivity in hMT+/V5 is anchored to an external, rather than somatotopic reference frame, which may be instrumental in constructing a stable representation of dynamic events for action despite continuous eye and hand movements (Burr & Morrone, 2011; Crespi et al., 2011; d’Avossa et al., 2007). To further delineate this role, we performed fROI-based MVPA within individually localized subregions of the right hMT+/V5, specifically MST and MT (Figures 5B and 5C). We found that tactile motion selectivity in an external frame of reference was evident in both subregions. The presence of shared crossmodal motion representations in both right MST and MT suggests that both subdivisions contribute to the alignment of multisensory motion information.

To identify regions beyond our a priori fROI hypotheses, we conducted group-level searchlight decoding analyses (Figure 4). Within-modality searchlight analyses confirmed that primary sensory cortices represent motion information specific to their respective modalities: decoding of visual motion directions (Figure 4B) revealed robust activity in the primary visual cortex (V1), hMT+/V5, and dorsal stream regions, while tactile motion decoding in both somatotopic (Figure 4A) and external (Figure 4C) reference frames uncovered extensive representations in the somatosensory cortices (S1 and S2), as well as hMT+/V5. Crucially, crossmodal searchlight decoding between vision and touch (Figures 4C and 4D) confirmed our fROI findings by showing that the right hMT+/V5 supports aligned visual and tactile motion representations within an external frame of reference. Beyond hMT+/V5, crossmodal alignment between visual motion and tactile motion (coded externally) also revealed additional regions in the frontal and parietal cortices, including the lateral intraparietal area (LIP), ventral intraparietal area (VIP) within the posterior parietal cortex (PPC), and premotor areas in the frontal cortex. These findings are consistent with prior work implicating the PPC and frontal regions in automatic remapping of touch into external spatial coordinates (Azañón & Soto-Faraco, 2008a; Bruns & Röder, 2010; Crollen, Albouy, et al., 2017; Soto-Faraco & Azañón, 2013), as well as in multisensory spatial integration (Andersen et al., 1997; Azañón & Soto-Faraco, 2008b; Lloyd et al., 2003; Serino et al., 2011). Together, our results demonstrate that hMT+/V5, along with a distributed fronto-parietal network, plays a central role in aligning motion direction representations across vision and touch.

It remains unclear whether the alignment of motion representations across the senses relies on a common, abstract frame of reference or instead involves remapping onto an anchor frame specific to a single modality (Avillac et al., 2005) (Pouget et al., 2002; Pouget & Sejnowski, 1997). An alternative possibility is that the fronto-parietal-hMT+/V5 network flexibly recruits different reference frames depending on task demands (Heed et al., 2015). For example, hMT+/V5 has been shown to encode motion in a non-retinotopic (external) frame of reference under low attentional load, but revert to a retinotopic frame of reference when attention is focused on the fovea during continuous and demanding visual discrimination tasks (Burr & Morrone, 2011; Crespi et al., 2011; d’Avossa et al., 2007; Gardner et al., 2008). Similarly, the posterior parietal cortex dynamically updates its coding, shifting from representing sensory information during target localization to representing movement goals during motor planning (Klautke et al., 2022). Our findings thus open the door to future investigations into how different nodes of the fronto-parietal-hMT+/V5 network flexibly cooperate to align crossmodal frames of reference in a task-dependent manner (Rohe et al., 2019; Rohe & Noppeney, 2015, 2016).

One potential alternative explanation is that imagining visual motion during tactile stimulation could evoke selective activity in hMT+/V5, allowing tactile motion directions to be decoded indirectly through visual imagery. Indeed, early visual areas such as V1 are known to be engaged during visual imagery (Kosslyn et al., 1993), and the direction of imagined visual motion can be decoded from V1 activity pattern (Emmerling et al., 2016). However, our crossmodal decoding analyses did not reveal aligned representations of visual and tactile motion directions in V1 (Figure S3). Thus, if the aligned motion representations we observed in hMT+/V5 were merely a byproduct of visual imagery, we would expect similar crossmodal effects to emerge in V1 — which was not the case. This suggests that the observed alignment in hMT+/V5 reflects genuine multisensory representation rather than imagery-related effects.

In conclusion, our study demonstrates that the right hMT+/V5, traditionally known for supporting visual motion signals, also represents tactile motion directions using an external frame of reference which aligns with the representation of visual motion directions. hMT+/V5 operates as part of a broader fronto-parietal network that supports shared, multimodal representations of motion signals, potentially enabling efficient and coherent perception of the dynamic external world. These findings open new avenues for investigating how biological and artificial neural systems implement coordinate transformations across sensory modalities to guide action in response to moving events in the environment.

## Acknowledgements

We would like to express our gratitude to Dr. Rémi Gau and Dr. Mohamed Rezk for their guidance on data management and analyses, and insightful discussions. fMRI data was acquired at the Cliniques Universitaires Saint-Luc (UCLouvain – Université Catholique de Louvain).

## Author contributions

Conception and design: IS, AM, OC

Participant Recruitment: IS

Data Collection: IS, CB, FC, AA

Data Analysis: IS analyzed data under the supervision of OC. CB provided support on data analysis.

Writing – original draft: IS, OC

Writing – review and editing: IS, CB, FC, AA, AM, OC

Funding Acquisition: AM, OC

Administrative support: IS, AM, OC

---

**Figure S 1.**
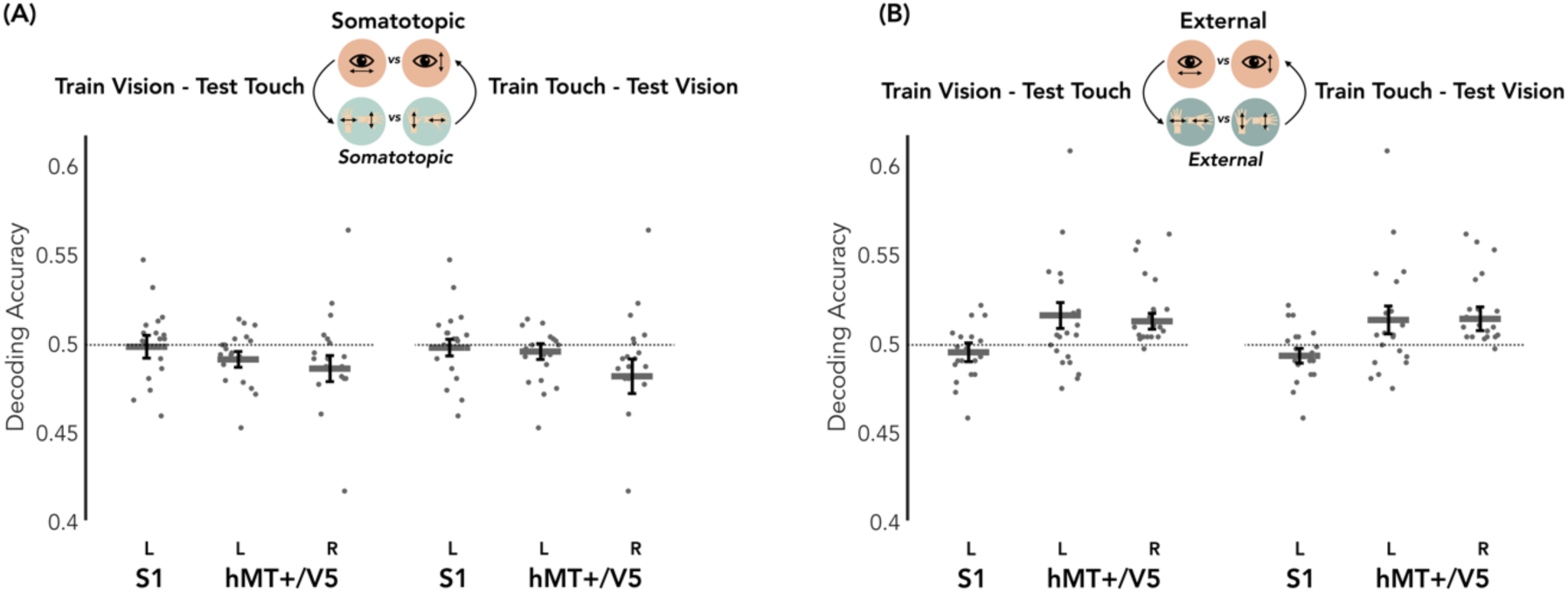
**Cross-Modal Decoding in the ROIs for different directions of train- and test-sets.** (A) Cross-modal classification between visual and tactile motion directions in the somatotopic frame of reference. The left side shows the results when the classifier is trained on visual modality and tested on tactile modality. The right side shows the results when the classifier is trained on tactile modality and tested on visual modality. (B) Cross-modal classification between visual and tactile motion directions in the external frame of reference. The left side shows the results when the classifier is trained on visual modality and tested on tactile modality. The right side shows the results when the classifier is trained on tactile modality and tested on visual modality.

**Figure S 2.**
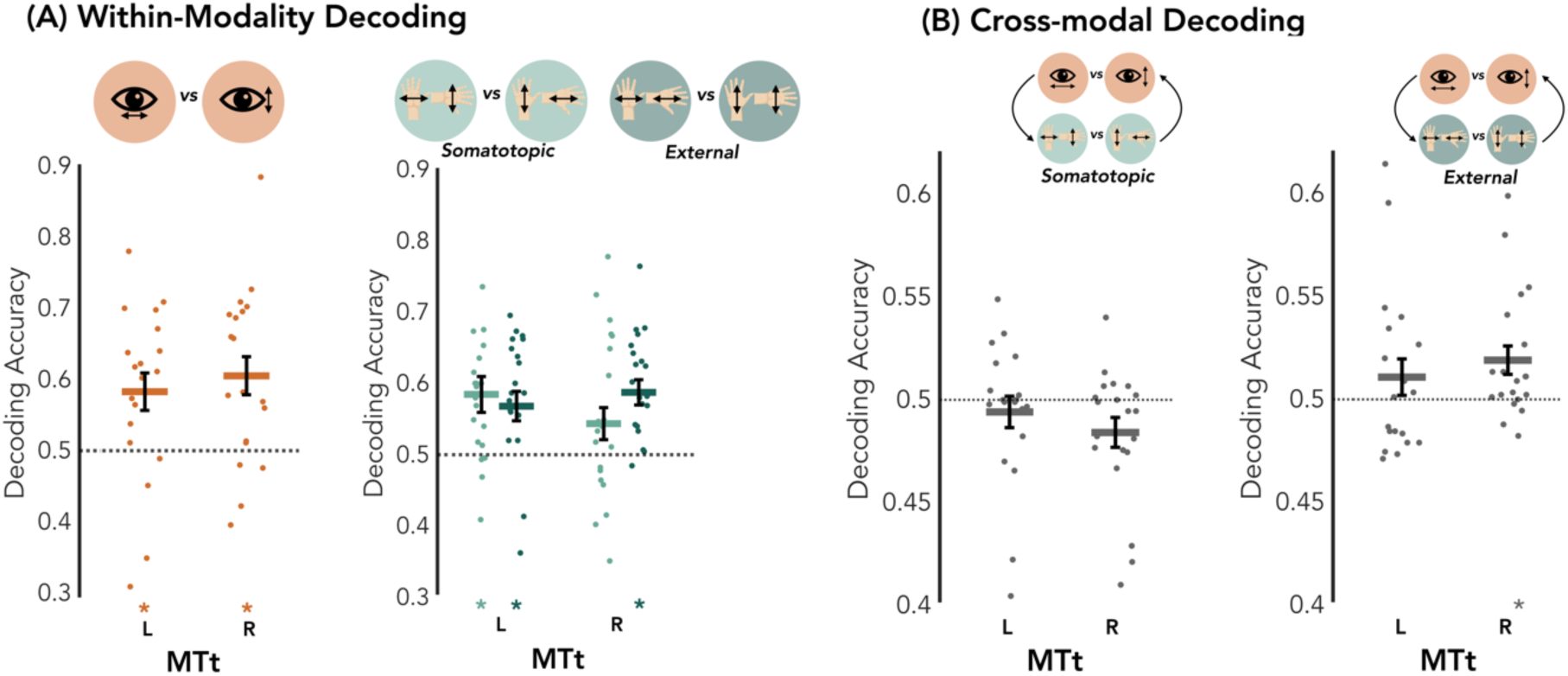
**Multivariate Pattern Analyses in the MTt.** (A) Within-Modality Decoding. Binary classification of visual (left panel) and tactile (right figure) motion directions across the two frames of references in each ROI. The y-axis represents the decoding accuracy, and the chance level is 0.5. (*p<0.05). (B) Cross-Modal Decoding. Cross-modal classification between visual and tactile motion directions across the two frames of reference – somatotopic (left panel) and external (right panel) within each ROI. In the left panel, the tactile motion directions are defined in somatotopic frame of reference and in the right panel, the tactile motion directions are defined in the external frame of reference.

**Figure S 3.**
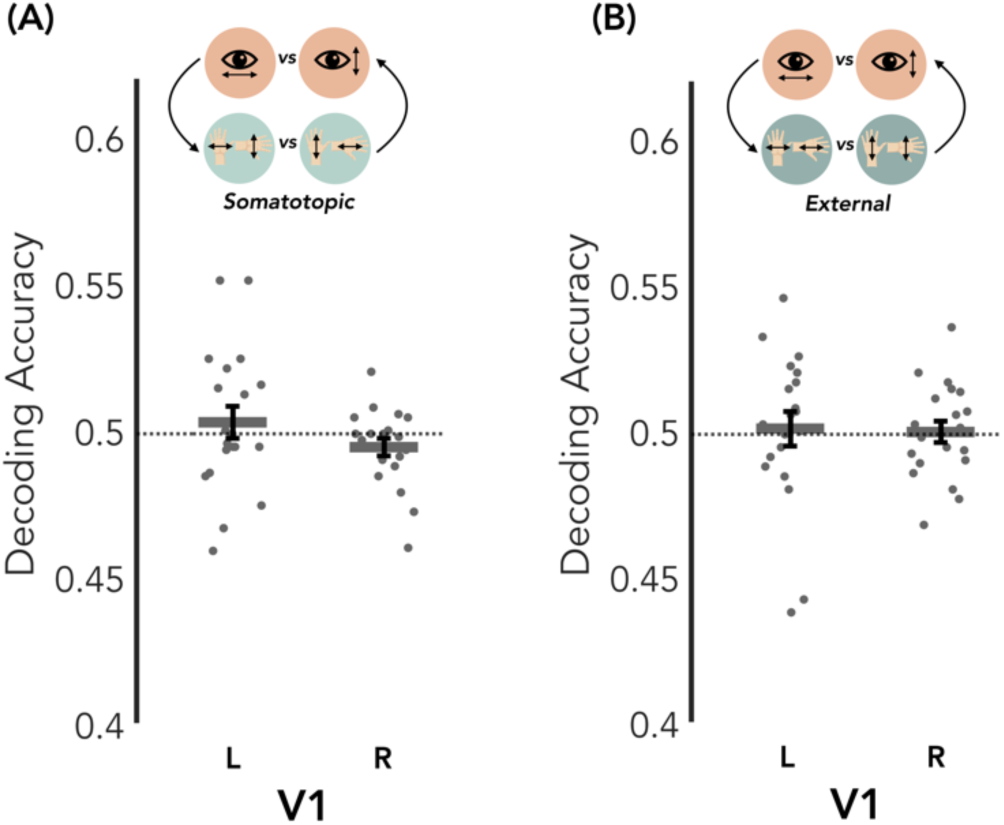
**Cross-Modal Decoding in V1.** Spherical fROIs (radius 8mm) of the visual area V1 were defined in each individual using visual motion localizer (motion > static). (A) Cross-modal classification between visual and tactile motion directions in the somatotopic frame of reference. (B) Cross-modal classification between visual and tactile motion directions in the external frame of reference.

**Table S 1.**
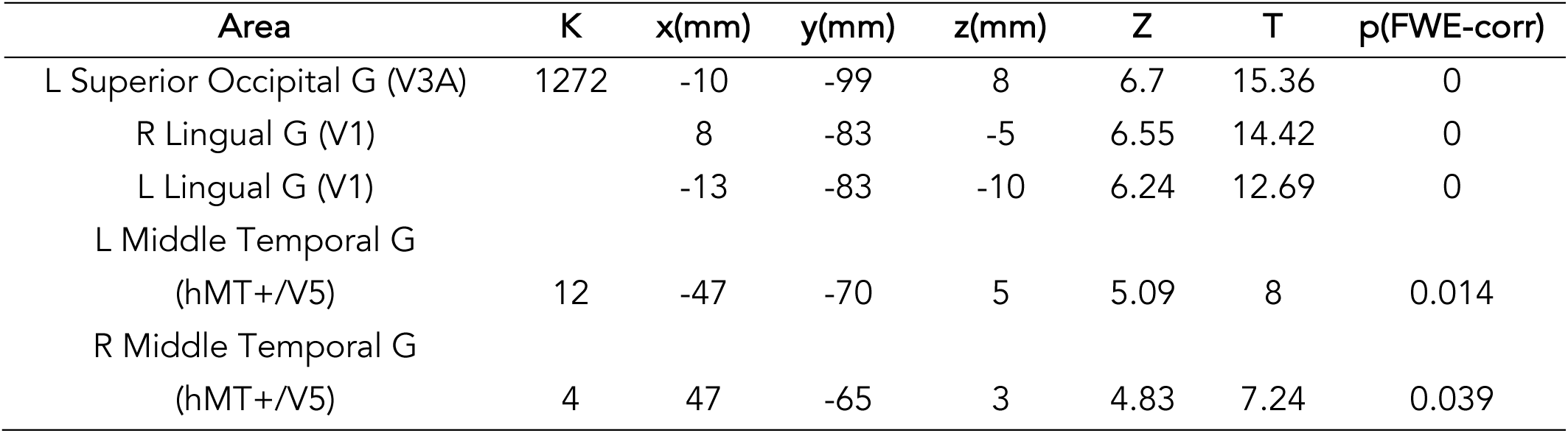
Whole-brain univariate analysis results from the visual motion localizer (visual motion > static). Results are corrected for multiple comparisons using family-wise error (FWE) correction across the whole brain at a significance level of p < 0.05. Coordinates are reported in MNI space, and regions are identified using the Automated Anatomical Labeling (AAL) atlas. L: Left; R: Right.

**Table S 2.** Whole-brain univariate analysis results from the tactile motion localizer. Results are corrected for multiple comparisons using family-wise error (FWE) correction across the whole brain at p < 0.05 or within small spherical volumes (*). Coordinates are reported in MNI space. Coordinates for small volume correction were derived from the independent visual motion localizer [MNI coordinates for left hMT+/V5: −47, −70, 5; right hMT+/V5: 47, −65, 3]. Regions are identified using the Automated Anatomical Labeling (AAL) atlas. L: Left; R: Right.

| Area | K | x(mm) | y(mm) | z(mm) | Z | T | p(FWE-corr) |
| --- | --- | --- | --- | --- | --- | --- | --- |
| L Postcentral G (S1) | 4312 | -55 | -21 | 52 | 5.29 | 8.65 | 0.005 |
| L Postcentral G |  | -44 | -26 | 63 | 5.19 | 8.33 | 0.007 |
| L Rolandic Oper (S2) |  | -42 | -16 | 13 | 4.97 | 7.66 | 0.017 |
| R Insula (S2) | 1149 | 39 | -8 | 11 | 4.32 | 5.95 | 0.18 |
| L Middle Temporal G<br>(hMT+/V5) | 104 | -52 | -70 | 8 | 3.19 | 3.8 | 0.026* |
| R Middle Temporal G<br>(hMT+/V5) | 122 | -52 | -42 | -10 | 2.98 | 3.47 | 0.043* |

**Table S 3.**
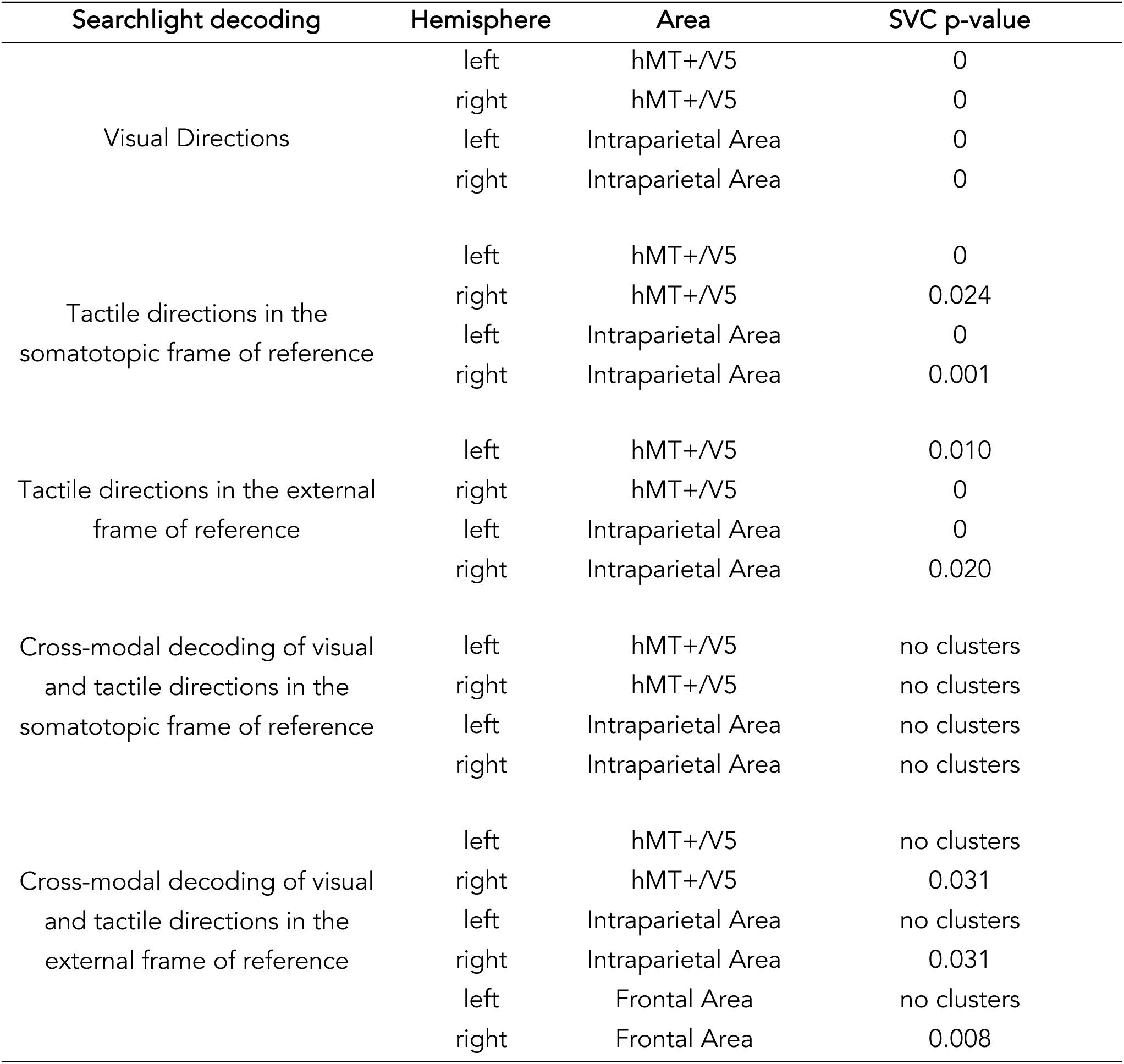
Summary Statistics of Searchlight Decoding. Small Volume Correction (SVC) was applied at the independently identified locations of left hMT+/V5 (−47, −70, 5 mm) and right hMT+/V5 (47, −65, 3 mm) from the visual motion localizer. Additionally, SVC was applied to parietal regions using coordinates from a prior study (Summers et al., 2009) that evaluated tactile motion selectivity [moving tactile > stationary tactile]: left Intraparietal Area (Talairach: −30, −50, 45; MNI: −30, −53, 49 mm) and right Intraparietal Area (Talairach: 42, −51, 49; MNI: 41, −55, 54 mm) and frontal regions using coordinates from a previous study (Lloyd et al., 2003) investigating the external representation of touch : left premotor cortex (−46, 8, 46 mm) and right premotor cortex (24, 4, 58 mm).

| Searchlight decoding | Hemisphere | Area | SVC p-value |
| --- | --- | --- | --- |
| Visual Directions | left | hMT+/V5 | 0 |
|  | right | hMT+/V5 | 0 |
|  | left | Intraparietal Area | 0 |
|  | right | Intraparietal Area | 0 |
| Tactile directions in the somatotopic frame of reference | left | hMT+/V5 | 0 |
|  | right | hMT+/V5 | 0.024 |
|  | left | Intraparietal Area | 0 |
|  | right | Intraparietal Area | 0.001 |
| Tactile directions in the external frame of reference | left | hMT+/V5 | 0.010 |
|  | right | hMT+/V5 | 0 |
|  | left | Intraparietal Area | 0 |
|  | right | Intraparietal Area | 0.020 |
| Cross-modal decoding of visual and tactile directions in the somatotopic frame of reference | left | hMT+/V5 | no clusters |
|  | right | hMT+/V5 | no clusters |
|  | left | Intraparietal Area | no clusters |
|  | right | Intraparietal Area | no clusters |
| Cross-modal decoding of visual and tactile directions in the external frame of reference | left | hMT+/V5 | no clusters |
|  | right | hMT+/V5 | 0.031 |
|  | left | Intraparietal Area | no clusters |
|  | right | Intraparietal Area | 0.031 |
|  | left | Frontal Area | no clusters |
|  | right | Frontal Area | 0.008 |

**Table S 4.** Behavioral performance across experimental conditions. N is the number of participants; mean is the mean performance accuracy calculated across participants; SD is the standard deviation; SEM is the standard error mean.

| Experiment | Condition | N | mean | SD | SEM |
| --- | --- | --- | --- | --- | --- |
| Visual Motion Localizer | Motion | 18 | 0.824 | 0.381 | 0.09 |
| Visual Motion Localizer | Static | 18 | 0.824 | 0.381 | 0.09 |
| Tactile Motion Localizer | Motion | 18 | 0.72 | 0.403 | 0.095 |
| Tactile Motion Localizer | Static | 18 | 0.731 | 0.409 | 0.096 |
| MT/MST Localizer | Motion-Left | 18 | 0.9 | 0.293 | 0.069 |
| MT/MST Localizer | Motion-Right | 18 | 0.861 | 0.318 | 0.075 |
| Visual | Vertical | 20 | 0.935 | 0.124 | 0.028 |
| Visual | Horizontal | 20 | 0.921 | 0.125 | 0.028 |
| Hand Up | Pinky-Thumb | 20 | 0.874 | 0.144 | 0.032 |
| Hand Up | Finger-Wrist | 20 | 0.863 | 0.149 | 0.033 |
| Hand Down | Pinky-Thumb | 20 | 0.864 | 0.187 | 0.042 |
| Hand Down | Finger-Wrist | 20 | 0.835 | 0.195 | 0.044 |

**Table S 5.** MNI Coordinates of the individually defined regions of interest (ROIs). Coordinates of bilateral hMT+/V5 were defined in each subject from the contrast [visual motion > visual static] in the visual motion localizer. Coordinates of left S1 were defined in each subject from the contrast [tactile motion > tactile static] in the tactile motion localizer.

|  | IS1 | lhMT+/V5 | rhMT+/V5 |
| --- | --- | --- | --- |
| S1 | -36.40, -28.80, 57.40 | -49.40, -57.40, 8.0 | 49.38, -64.79, 5.37 |
| S2 | -44.33, -28.09, 54.98 | -41.57, -58.37, 12.72 | 46.63, -61.12, 5.37 |
| S3 | -42.49, -21.67, 58.66 | -46.17, -64.79, 3.53 | 46.80, -62.60, 0.20 |
| S4 | -42.49, -30.85, 55.90 | -43.41, -66.63, 9.04 | 46.63, -60.21, 2.61 |
| S5 | -39.74, -26.26, 58.66 | -46.17, -60.21, 5.37 | 45.71, -58.37, 6.29 |
| S6 | -39.0, -26.20, 62.60 | -44.33, -64.79, 2.61 | 43.87, -61.12, -0.15 |
| S7 | -44.33, -19.84, 56.82 | -46.80, -67.80, 2.80 | 48.47, -62.04, 0.20 |
| S8 | -41.60, -24.42, 58.66 | -46.17, -66.63, 0.20 | 46.80, -60.21, 4.45 |
| S9 | -38.90, -28.09, 58.66 | -44.33, -66.63, 8.12 | 45.71, -62.04, 0.20 |
| S10 | -37.90, -28.09, 58.66 | -44.33, -66.63, 5.37 | 45.71, -62.04, 2.61 |
| S11 | -39.0, -28.09, 56.82 | x | x |
| S12 | -42.49, -24.42, 56.82 | -48.01, -67.55, 5.40 | x |
| S13 | -42.49, -23.60, 59.57 | -44.33, -64.79, -0.15 | 49.40, -61.12, -2.90 |
| S14 | -44.33, -28.09, 55.90 | -42.49, -66.63, 1.69 | 46.80, -62.96, -1.06 |
| S15 | -43.41, -22.59, 54.98 | -46.80, -65.20, 0.20 | 46.63, -60.21, 2.61 |
| S16 | -44.20, -31.40, 49.60 | -44.20, -67.80, 0.20 | 45.71, -62.04, -0.15 |
| S17 | x | x | x |
| S18 | x | x | x |
| S19 | -39.0, -28.09, 56.82 | -49.40, -62.60, 2.80 | 54.60, -57.40, 0.20 |
| S20 | -35.14, -28.09, 59.57 | -49.40, -65.20, 2.80 | 45.71, -62.96, 3.53 |

**Table S 6.** Voxel counts in the right hMT+/V5 complex and its subregions – MST and MT. Number of voxels within each region of interest (ROI): right hemisphere hMT+/V5, MST, and MT. X denotes participants not tested for MT/MST visual motion localizer. – denotes that there were no clusters observed.

| Subject | Right<br>hMT+/V5 | Right<br>MST | Right<br>MT |
| --- | --- | --- | --- |
| S1 | 192 | 16 | 176 |
| S2 | 312 | 133 | 179 |
| S3 | 243 | 10 | 233 |
| S4 | 306 | 212 | 94 |
| S5 | 190 | 21 | 169 |
| S6 | 270 | 112 | 158 |
| S7 | 208 | 10 | 198 |
| S8 | 305 | 40 | 265 |
| S9 | 298 | 52 | 246 |
| S10 | 301 | 134 | 167 |
| S11 | - | - | - |
| S12 | - | - | - |
| S13 | 164 | - | - |
| S14 | 367 | 34 | 333 |
| S15 | 440 | 110 | 330 |
| S16 | 344 | 4 | 340 |
| S17 | x | x | x |
| S18 | x | x | x |
| S19 | 58 | 21 | 37 |
| S20 | 379 | 11 | 368 |

**Table S 7.** Dice Coefficient between the individually localized hMT+/V5 and MTt. The dice coefficient depicts the spatial overlap between individually localized hMT+/V5 and MT regions within each participant.

| Subject | Left Hemisphere | Right Hemisphere |
| --- | --- | --- |
| S1 | 0.669354839 | 0.798319328 |
| S2 | 0.065306122 | 0.359832636 |
| S3 | 0.186234818 | 0.260162602 |
| S4 | 0.21399177 | 0.333333333 |
| S5 | 0 | 0.348178138 |
| S6 | 0.551440329 | 0.123966942 |
| S7 | 0.048780488 | 0.11814346 |
| S8 | 0.721311475 | 0.533333333 |
| S9 | 0.202531646 | 0.271604938 |
| S10 | 0.878661088 | 0.685714286 |
| S11 | 0.596774194 | 0.205761317 |
| S12 | 0.468619247 | 0.205761317 |
| S13 | 0.040983607 | 0.226720648 |
| S14 | 0.04 | 0.193277311 |
| S15 | 0.227642276 | 0.216666667 |
| S16 | 0.696721311 | 0.721311475 |
| S17 | 0.596774194 | 0.205761317 |
| S18 | 0.596774194 | 0.205761317 |
| S19 | 0.596774194 | 0.205761317 |
| S20 | 0.308943089 | 0.545454545 |

